# Commensal microbiome dysbiosis elicits IL-8 signaling to drive fibrotic skin disease

**DOI:** 10.1101/2023.09.19.558395

**Authors:** Wenyu Zhang, Qili Peng, Xian Huang, Qing Huang, Zhiliang Zhang, Fuli Li, Naisheng Zheng, Binsheng Shi, Zhihong Fan, Tomasz Maj, Rui Chen

## Abstract

Commensal bacteria are core players in wound healing whose function in the opposite pathophysiological process-scarring is presently unclear. Here, we document the association between bacteria and a specific skin fibrotic disease-keloid, which might offer a promising avenue for translational practice.

**ABSTRACT:** Wound healing is an intensely studied topic involved in many relevant pathophysiological processes, including fibrosis. Despite the large interest in fibrosis, the network that related to commensal microbiota and skin fibrosis remain mysterious. Here, we pay attention to keloid, a classical yet intractable skin fibrotic disease to establish the association between commensal microbiota to scaring tissue. Our histological data reveal the presence of microbiota in the keloids. 16S rRNA sequencing characterize microbial composition and divergence between the pathological and normal skin tissue. Moreover, the data show elevation of interleukin-8 both in the circulation and keloid tissue, which elicited the collagen accumulation and migratory program of dermal fibroblasts via CXCR1/2 receptor. Our research provides insights into the pathology of human fibrotic diseases, advocating commensal bacteria and IL-8 signaling as useful targets in future interventions of recurrent keloid disease.

## INTRODUCTION

Wound healing in adult human tissues generally involves fibrotic scar instead of tissue regeneration, which impair quality of life and even cause death. As a mirror to other organs’ wound healing, fibrotic skin diseases of great interests, while the keloid is an always overlooked one (1). Keloids are thick, raised scars that grow beyond the original wound area (2, 3). The scars may not be deadly, but they cause chronic itching, pain, and cosmetic disfigurement, leading to a poor quality of life, anxiety, and depression (4). Keloid is refractory to current treatment regimen, including surgery, corticosteroid medication, and radiation (3). Therefore, it is an urgent clinical need to complete understand its complex pathogenesis. The basis of keloid formation is fibroblasts hyperproliferation and deposition of large amounts of collagen type I and III (2, 3). The initiation and support of this fibroblastic pathology remains elusive. Previous research indicates that keloids are related to genetic features (5, 6), mechanical properties of the skin (7), nutrition and metabolism (8), composition of local hematopoietic compartment (9–11), and neuro-endocrine interactions (12). These different factors contribute to keloid formation, but no single aspect is the main cause.

Emerging evidences show links between the microbiota and the development of fibrosis. For example, intestinal microbiota is a key driver of inflammatory bowel disease, which develops further to intestinal fibrosis (13). Also, gut bacteria are linked with liver fibrosis in non-alcoholic fatty liver disease (14). Lung microbiota elicits interleukin-17B signaling pathway, which promotes the progression of pulmonary fibrosis (15). In a mouse model, genetic or chemical inhibition of Toll-like receptor 4 (TLR4), essential sensor of bacterial products, significantly reduces fibrosis progression (16, 17). These studies suggest that the disturbance of the microbial community has an impact on profibrotic events in different organs. Therefore, it is reasonable to ask whether and how microbiota contribute to fibrosis of the skin.

Skin is one of the critical barrier tissues directly interacting with microbiota. The presence of microorganisms was well studied in dermal infections for decades. Recent sequencing methods show that skin is a complex ecological network of microorganisms and the host (18). It is also well established that both commensal and pathogenic microbiota affect the outcome of wound healing (18, 19). However, to our knowledge, the microbiota was not studied in keloids. In this paper, we used 16S rRNA gene sequencing of hospitalized patients to gain a landscape of the microbial community. We show a significant dysbiosis of cutaneous microbiome between keloid lesion and non-keloid skin. Next, we analyzed potential cytokines and chemokines. We demonstrate that interleukin-8 (IL-8), also known as CXCL8, is elevated in keloids, which is also supported by clinical tests for circulating cytokines. We also checked the spatial distribution of IL-8 in the keloid microenvironment. Next, we defined the role of IL-8 on phenotypic and functional regulation of dermal fibroblasts. We show that IL-8 could promote profibrotic events, including migration, collagen deposition, and contraction of dermal fibroblasts. The findings provide new insights into etiology of keloid disease and identify signals helping the proper tissue repair. Thus, it reveals potential new therapeutic avenues to optimize clinical management of the disease.

## RESULTS

### Microbiota colonize keloids

To check the possibility that bacteria reside in the keloid tissues, we performed the general immunohistochemical (IHC) staining of lipoteichoic acid (LTA) and lipopolysaccharide (LPS) in clinical samples. These molecules are distinguishing features of Gram-positive and Gram-negative bacteria, respectively. Both locally or systemically, they induce immune-related signaling as pathogen-associated molecular patterns (20–22). According to the staining, we detect LTA mainly in epidermis, but not in the deeper parts of the tissue (Figure 1A and Supplemental Figure 1A). In contrast, LPS is distributed both in epidermis and dermis (Figure 1A and Supplemental Figure 1A). These data were further characterized by detailed histological scoring. In the keloid tissue, LPS displays similar scoring values in epidermis and dermis (Figure 1B and Supplemental Figure 1B), while LTA is distributed differentially, located mainly in epidermis (Figure 1C). Interestingly, the IHC score for LPS is higher both in epidermis and dermis of keloids compared to normal skin (Figure 1D), while LTA shows higher concentration in keloid than normal epidermis, but not dermis (Figure 1E). It implies that microbial components in healthy and keloid skin are different.

**Figure 1.**
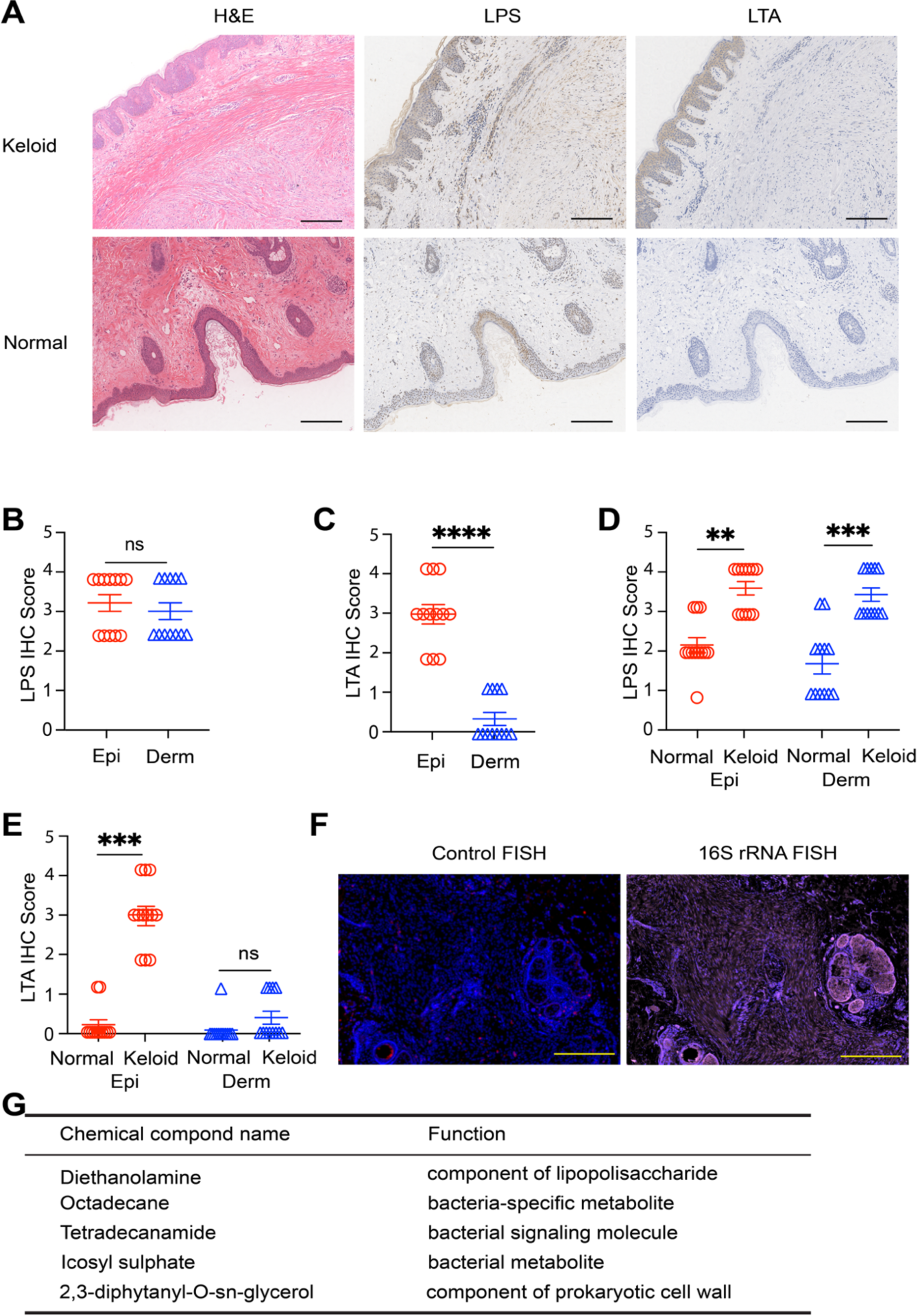
Colonization of keloid and normal skin by bacteria. (**A**) Representative pictures of histological staining of keloid in normal tissue (left column), and IHC staining specific for LPS and LTA. Scale bars, 200 μm. (**B** and **C**) Difference in IHC scoring for LPS (B) and LTA (C); (**D** and **E**) Comparison of IHC scoring values between epidermis and dermis in normal and keloid skin tissues for LPS (D) and LTA (E). (**F**) Representative picture of 16S rRNA FISH, depicting specific staining (right) and control staining with not specific oligonucleotides (left). n = 6, scale bars, 500 μm. (**G**) The list of bacteria-related chemicals identified in the keloid tissues by non-targeted metabolomic analysis (n = 6). Statistical significance (Wilcoxon test): \*\**p*<0.01, \*\*\**p*<0.001 and \*\*\*\**p*<0.0001, respectively; ns, not significant.

Though IHC data may suggest that keloids are colonized mainly by Gram-negative bacteria (Supplemental Figure 1C), the retention of bacterial products in the tissues can last days or weeks. Therefore, they may be not representative of general bacteria configuration (23, 24). We conducted fluorescent in situ hybridization (FISH) to directly detect bacterial 16S rRNA gene and confirm the existence of bacteria. Similar to IHC, we observed the widespread presence of bacterial rRNA in the keloid tissue (Figure 1F), which was also higher compared to normal skin (Supplemental Figure 1D). We also performed the non-target metabolomic analysis for keloid tissues (Figure 1G). We detected some microbiota-associated components, implying the presence of microorganisms. Importantly, 16S rRNA is also linked to immunostaining of fibroblast activation protein alpha (FAP), the marker of fibroblasts in keloids (25) (Supplemental Figure 1E). Together, these data show that keloid tissues contain bacteria different from normal skin.

### Keloid and healthy skin exhibit similar surface microbial community

Possible bacterial dysbiosis in the epidermis, and later in the dermis, may result from a different composition of microbiome on epidermal tissues. As a scar tissue, keloids have a different composition of glands and hair follicles. The surface of skin in these areas shows different chemical composition and may support various bacteria that further infect the keloid scar. To test this point of view, we performed a swab test, collecting microbiota from the surface of keloid scar and adjacent healthy skin of patients. The material was further studied by 16S rRNA sequencing and bioinformatic analysis. The microbiota composition on normal skin and keloids was similar but varied between specimens. The four most abundant phyla were *Proteobacteria*, *Bacteroidota*, *Firmicutes* and *Actinobacteria* (Figure 2A), typical groups identified in human skin. Both on normal skin and keloids, the most predominant genus was *Cutibacterium*, followed by *Staphylococcus*, *Corynebacterium*, and *Acinetobacter* (Supplemental Figure 2, A and B). Importantly, from over 300 identified amplicon sequence variants (ASV), 80% were shared between healthy and keloid skin, 11% were specific for normal skin, and 9% for keloids (Figure 2B). This result is also confirmed by alpha diversity values. They are slightly lower in keloids, whereas not statistically different (Figure 2C). Further, the graph permutation analysis show that the distance among normal skin and keloids swab specimens are mixed with high level of similarities (Figure 2D). That is also confirmed by the small beta diversity of the samples (Figure 2E). We did not find any significant results in the metagenomic functional analysis based on KEGG pathways, except for four pathways with p-values below 0.01 (Supplemental Figure 2C). The composition of microbial communities is almost the same in keloids and normal skin on the tissue surface.

**Figure 2.**
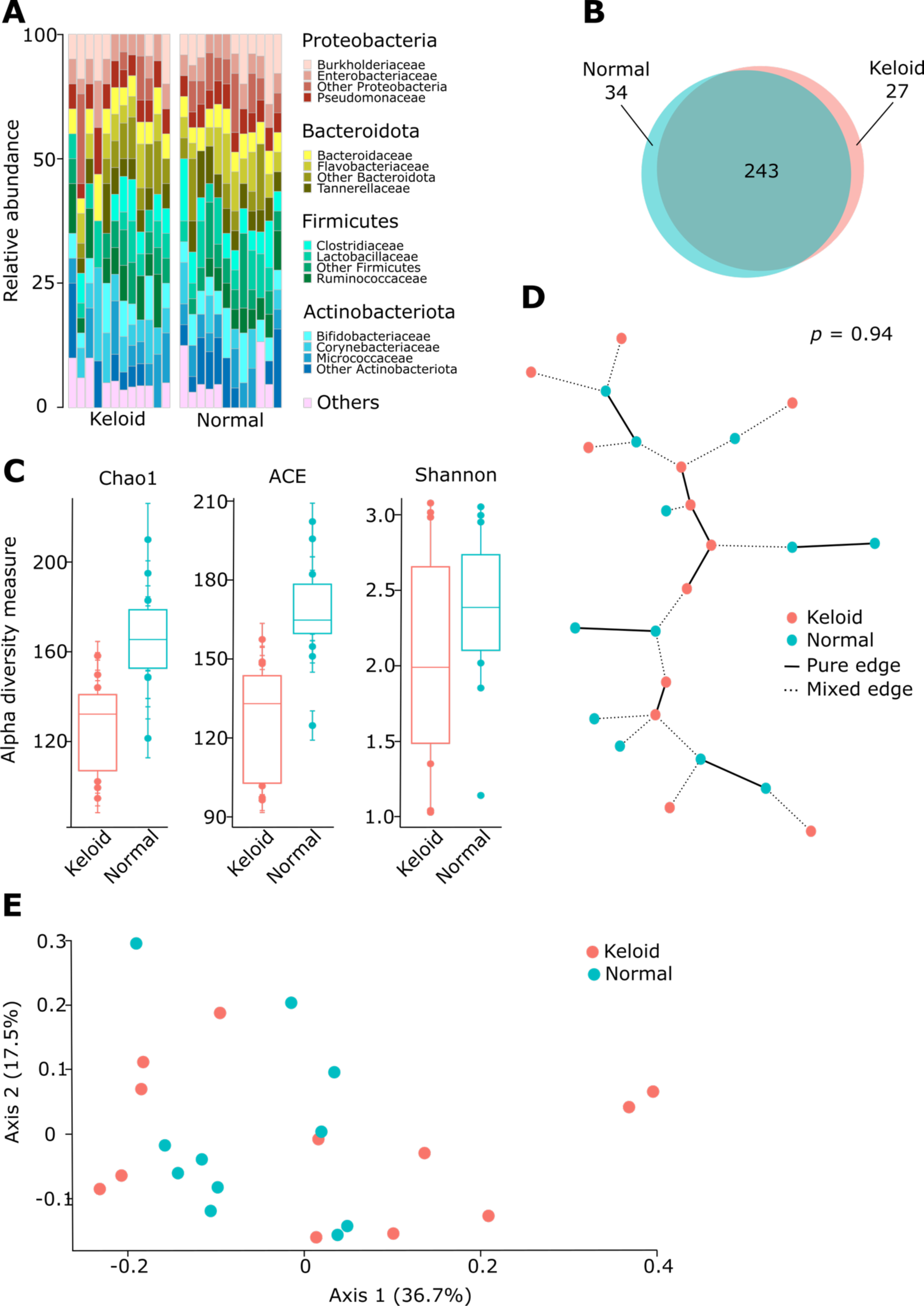
Composition of bacteria on skin surface of normal and keloid tissue. The swab samples were taken from keloids and normal skin (n = 12 in each group) and subjected to 16S rRNA sequencing and analysis. (**A**) Composition of bacteria at the level of phylum and class. The graph shows four the most abundant phyla, while all other are annotaded as “Others”. In similar manner, each phylum also shows three the most abundant classes. Each column shows separate keloid or normal sample. (**B**) Venn diagram representing the number of taxa specific for each type of sample and shared between samples. (**C**) The differences in alpha diversity measured by three different methods. (**D**) Graph showing differences between composition of bacteria residing on the surface of keloids or healthy skin. (**E**) Beta diversity analysis of swab assays between taken from healthy and pathological skin surface.

### Deep bacterial communities in healthy skin and keloids are different

As shown before (Figure 1), keloid samples compared to normal skin reveal potentially higher number of bacteria. Following this, we characterized bacterial composition also inside the keloid and healthy skin tissues. We collected samples from five keloid patients and compared the central part of the keloid with the marginal, healthy-looking adjacent skin. We found that in all cases, the abundance of bacteria was higher in keloid tissue than in the healthy part, though the differences are variable (Figure 3A). In contrast to the surface microbiota, the internal bacteria are different between keloid and normal tissues at the level of phylum and class (Figure 3B). The differences between samples within keloid and adjacent tissues are more pronounced. In all cases, similar to the surface bacteria, the predominant phyla are *Proteobacteria*, *Bacteroidota*, *Firmicutes* and *Actinobacteria*. However, the alpha diversity of central keloid samples is significantly higher than normal skin (Figure 3C), implying the dysfunction of the barrier. The surface composition of microbiota had twice as many ASVs as the swab tests with the same filtering method and settings. Both tissues share only 32% of ASVs, and 59% of detected taxons were keloid-specific (Figure 3D and Supplemental Figure 3A). In terms of beta-diversity, the samples separate well (Figure 3E) and the differences are statistically significant (Supplemental Figure 3B). Finally, the metagenomic functional analysis shows 892 different metabolic functional pathways (*p* < 0.001) that are clearly divided between bacteria isolated from keloid and healthy skin, and form clusters of healthy and keloid tissues (Figure 3F). The data indicates that bacterial phenome of keloid tissue display more bacteria, more diversity, and different compositions compared to the surface and normal skin microbiome.

**Figure 3.**
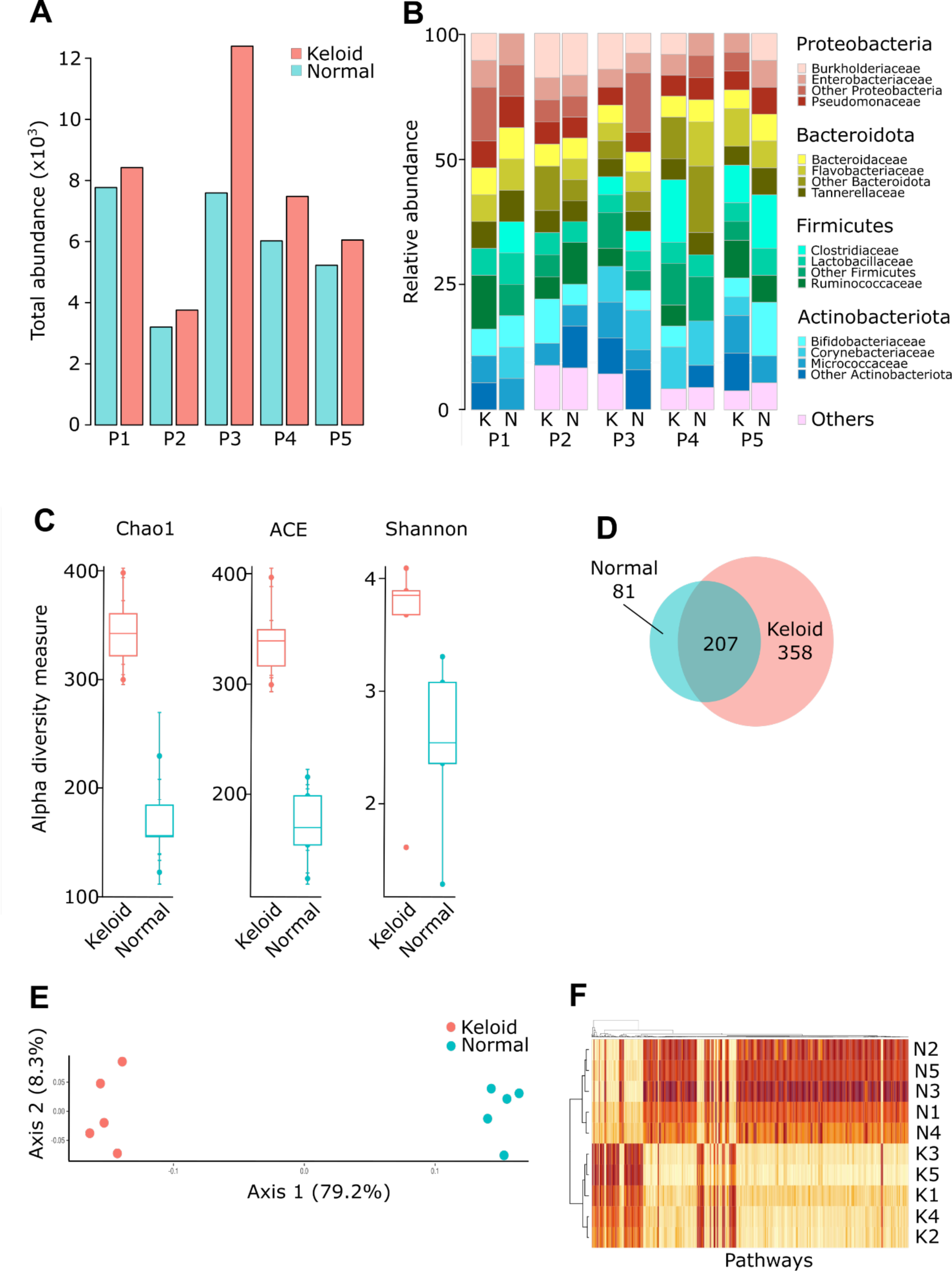
Bacterial community inside normal and pathological skin samples. Normal adjacent skin and central keloid tissues (n = 5 per group) were sampled from keloid-bearing patients, and subjected to 16S rRNA analysis. (**A**) Total abundance of bacteria in adjacent and pathological tissue. P1-P5, patient 1 to patient 5. (**B**) Comparison of bacterial composition in keloid and adjacent healthy skin samples at phylum and class level. K, keloid; N, normal tissue; P1-P5, patient identifier. (**C**) Comparison of alpha diversity between normal and keloid tissues. (**D**) Venn diagram showing the specific and overlapped taxa number in normal and keloid microbial profiling. (**E**) Analysis of beta diversity performed by PCA-wunifrac method. (**F**) Heatmap representing metagenomic analysis of microbiota colonizing normal and keloid samples. The data show functional pathways analyzed with tax4FUN method with \*\*\**p*<0.001.

### Overproduction of IL-8 in keloids

Keloids develop a specific inflammatory environment that supports fibrosis. Therefore, we try to assess whether we can detect cytokines related to anti-bacterial immune responses. First, we tested blood from keloid patients. At the plasma level, we screen several cytokines known as pro- (Figure 4A and Supplemental Figure 4A) or anti-inflammatory (Supplemental Figure 4B) factors. Some of them, like IL-1β or IL-5 are not detected in keloid patients, and others are present in range between 20 to 40% patients. Interestingly, IL-8 is one of the top 3 cytokines associated with keloids (Figure 4A). This cytokine/chemokine is critical in fast response to bacteria, and fibroblasts, the key players in keloid development, also express the receptors for IL-8 (26, 27). Having this, we studied the co-localization of IL-8 protein with fibroblast and macrophage populations. We used α-SMA as a marker of fibroblasts and CD68 as a marker of macrophages. Though macrophages are considered as a potent IL-8 producers, we observed that α-SMA^+^ cells are the main sources of IL-8 in keloids (approximately 40% of cells on average, Figure 4, B and C). They are also the most closely located to IL-8 in various distances (Figure 4D). Since we lack animal models of keloids, we used *in vitro* assay to link bacteria with IL-8 keloid production. We treat the fibroblasts with heat-killed *Salmonella typhimurium* or *Staphylococcus aureus*, as examples of Gram-negative and Gram-positive bacterial strains. In both cases, the addition of bacteria induced the production of IL-8 (Figure 4, E and F). The data show that bacteria-activated fibroblasts produce IL-8 protein in the keloid environment.

**Figure 4.**
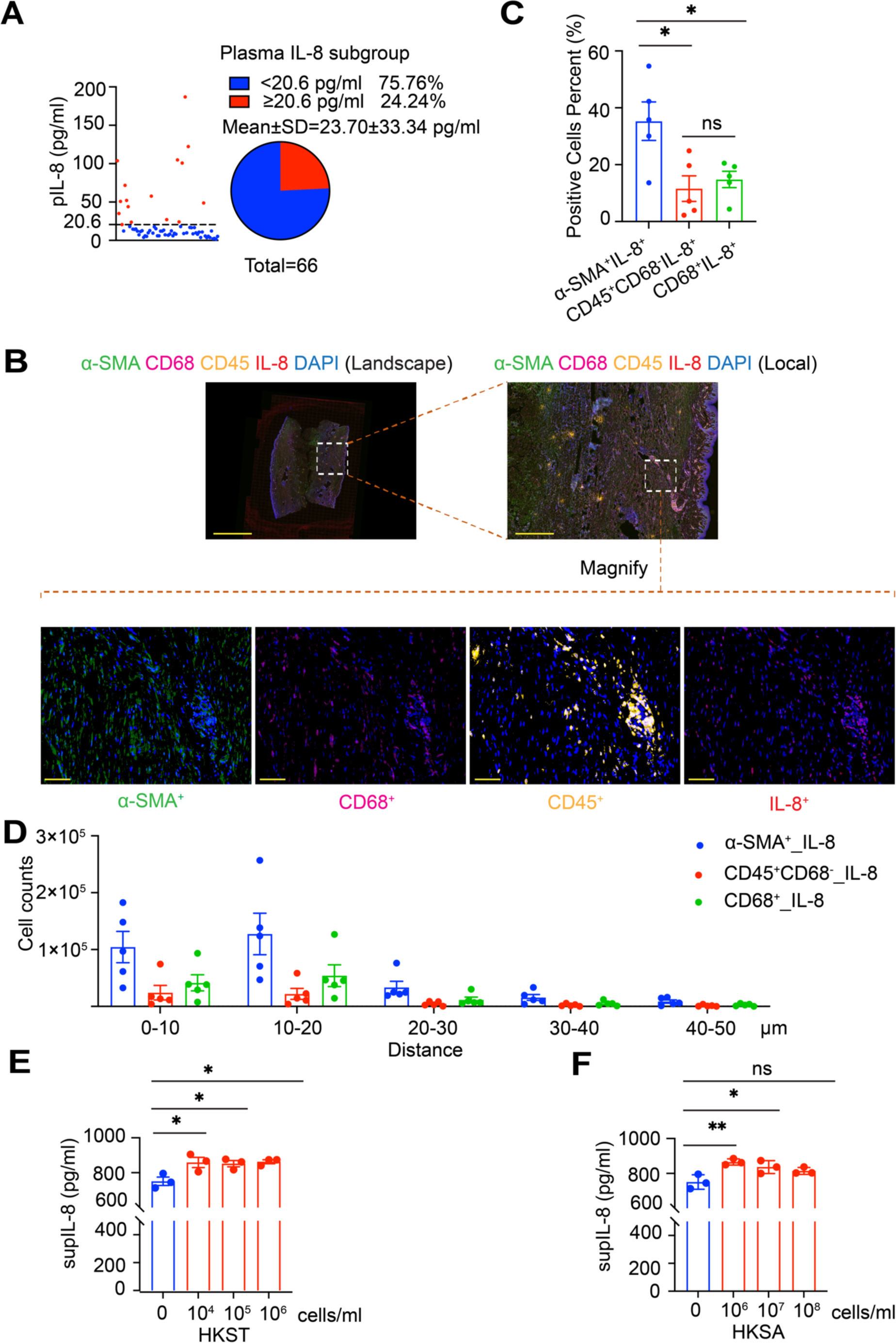
Fibroblasts are the feasible source of IL-8 in keloids. (**A**) Percentage of patients in tested IL-8 chemokine subgroup in circulation. Reference values (RVs) are as below: IL-8=0∼20.60 pg/mL, The number of patients (n=66) is indicated. (**B**) Representative fluorescence image of samples from 5 patients showing the simultaneous detection of α-SMA (green), CD68 (magenta), CD45 (SpGold), IL-8 (CY5, red) and 4’,6-diamidino-2-phenylindole (DAPI; blue) in human keloid. Scale bars, 100 μm. Magnification is 40x. (**C**) Statistic analysis of indicated cell percent based on mIHC staining data from (B). Mean ± SEM. ^ns^*p*>0.05; \**p*<0.05. Normality and Lognormality Tests followed by two-tailed unpaired Student’s t test. (**D**) Cell proximity analysis of indicated cells in the given distance based on mIHC staining data from (B). Quantitative immunofluorescence was processed by HALO software. (**E**) Dermal fibroblasts were treated with variable dosages of Heat Killed *Salmonella typhimurium* (HKST) for 24h. IL-8 concentration in the supernatant was measured by ELISA assays, n = 3 per subgroup. Normality and Lognormality Tests followed by one-way ANOVA was performed for the statistical analysis. Mean ± SEM. *p < 0.05. (**F**) Dermal fibroblasts were treated with variable dosages of heat-killed *Staphylococcus aureus* (HKSA) for 24h. IL-8 concentration in the supernatant was measured by ELISA assays, n = 3 per subgroup. Normality and log-normality tests followed by one-way ANOVA was performed for the statistical analysis. Mean ± SEM. ^ns^*p*>0.05; \**p*<0.05; \*\**p*<0.01.

### IL-8 mediates the changes in skin fibroblast behavior

To further determine the role of IL-8 on wound healing, we checked IL-8 effect on fibroblast’s proliferation and migration. Cell counting assays and FACS analysis on CFSE-labeled primary fibroblasts upon IL-8 stimulation showed no significant effect on the proliferation (Figure 5A and Supplemental Figure 5A). Our results of a trans-well assay displayed increased fibroblast migration when treated with recombinant human IL-8 (rhIL-8) (Figure 5, B and C). Also, we performed a separate scratch assay to confirm the effect of rhIL-8 treatment. Administration of rhIL-8 increased the ability of fibroblasts to migrate and seal the in vitro “wound” (Supplemental Figure 5, B and C). These data indicate that IL-8 increases the migration of skin fibroblasts.

**Figure 5.**
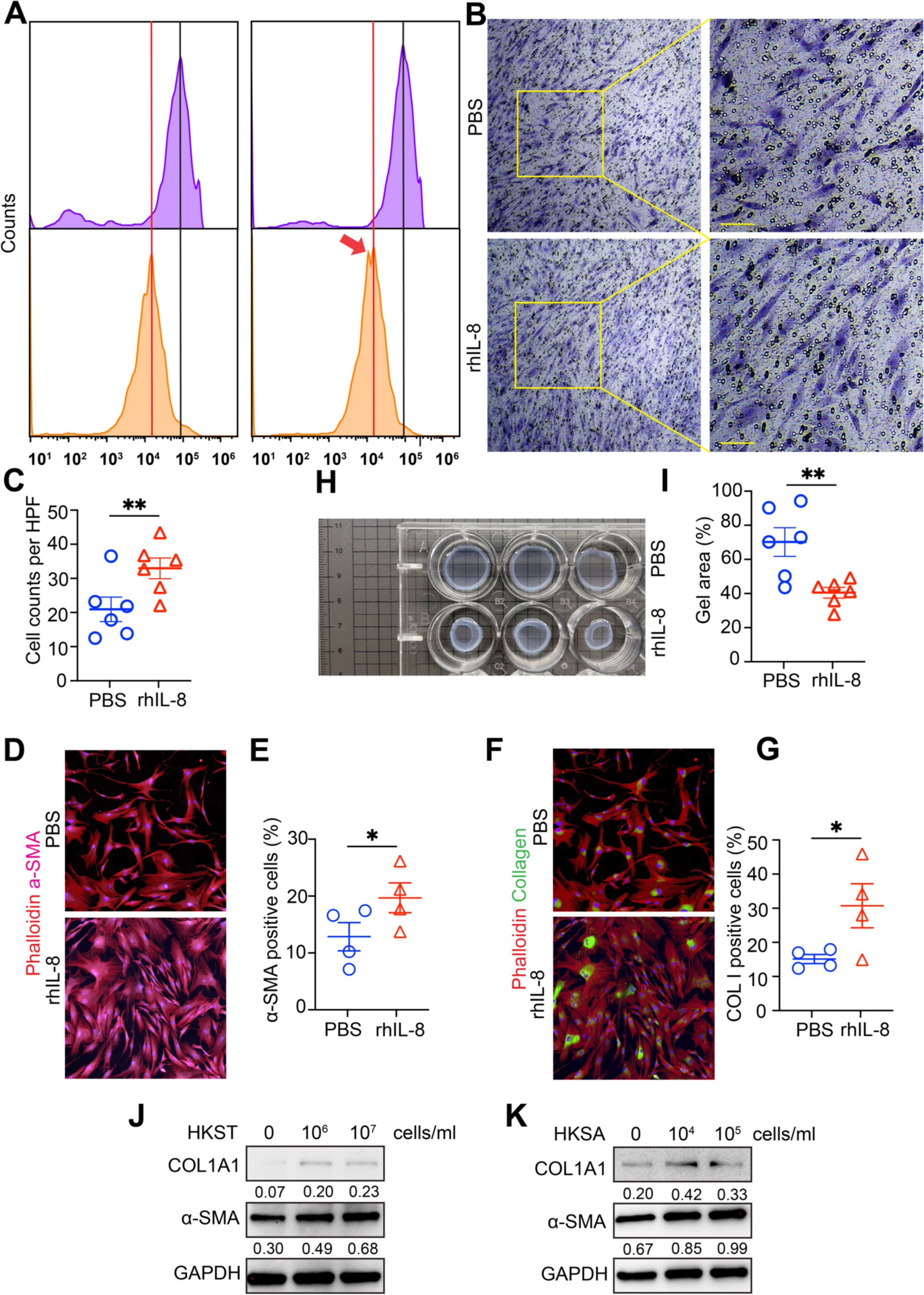
The Role of IL-8 on phenotype and functionality of dermal fibroblast. (**A**) FACS analysis on proliferation of CFSE-labeled dermal fibroblasts in response to vehicle (PBS) and rhIL-8 (50 ng/mL) (representative of two assays). (**B**) Representative images of trans-well assays on vehicle (PBS) and IL-8 (50 ng/mL) treated samples (n = 3 technical replicates, representative of two assays). (**C**) Statistic analysis of cell number based on photographed images from (B). Mean ± SEM. **p < 0.01. Normality and log-normality tests followed by two-tailed unpaired Student’s t test. HPF (high-power field). (**D**) Representative fluorescence images of independent samples from patients (n = 4) showing the detection of α-SMA (magenta). Phalloidin (red) was used for reflecting cell morphology. Scale bars, 200 μm. (**E**) Statistic analysis of α-SMA positive cell percent based on photographed images from (D). Mean ± SEM. \**p*<0.05. Normality and log-normality tests followed by two-tailed unpaired Student’s t-test. (**F**) Representative fluorescence images of independent samples from patients (n = 4) showing the detection of collagen (green). Phalloidin (red) was used for reflecting cell morphology. Scale bars, 200 μm. (**G**) Statistic analysis of collagen positive cell percent based on photographed images from (E). Mean ± SEM. \**p*<0.05. Normality and log-normality tests followed by two-tailed unpaired Student’s t-test. (**H**) Gel contraction analysis of dermal fibroblasts with indicated treatment (n = 3 technical replicates, representative of two assays). (**I**) Gel area (%) were calculated and analyzed based on the data from (H). Mean ± SEM. \*\**p*<0.01. Normality and log-normality tests followed by two-tailed unpaired Student’s t-test. (**J**) Dermal fibroblasts were treated with variable dosages of heat-killed *Salmonella typhimurium* (HKST) for 24 h. Indicated proteins were detected by western blot. 1 of 2 western blots is shown. (**K**) Dermal fibroblasts were treated with variable dosages of heat-killed *Staphylococcus aureus* (HKSA) for 24h. Indicated proteins were detected by western blot. Gray values were calculated by ImageJ, numerical values were calculated as a density of a given band to the density of GAPDH band ratio. 1 of 2 western blots is shown.

Myofibroblasts, a distinctive subset of activated fibroblasts has been implicated to be essential cells for extracellular matrix production, which is identified by a cardinal marker, α-SMA (28, 29). We initially found the elevated expression of α-SMA in response to IL-8 treatment (Figure 5, D and E). We therefore turned to check the production of collagen, a main component of the extracellular matrix. Treatment with IL-8 promoted the production of collagen I (Figure 5, F and G). Moreover, we performed a gel contraction assay to test contractile properties of fibroblasts upon IL-8 treatment. The result showed enhanced contractility of IL-8 treated fibroblasts (Figure 5, H and I). By contrast, neutralizing antibody of IL-8, CXCR1 inhibitor reparixin, or CXCR2 inhibitor (SB225002) hindered the contractility of fibroblasts in the presence of IL-8 (Supplemental Figure 5, D and E). In addition, we showed microbial cues could also induced fibroblast-myofibroblast transformation, as evidenced by elevated expression of α-SMA and collagen I (Figure 5, J and K, and Supplemental Figure 5, F and G), suggesting a plausible link. These data suggest that IL-8 signaling indeed mediates skin fibroblast activation.

Next, an RNA-seq was performed to explore deeper of the role of IL-8 on skin fibroblasts. We observed some ribosomal proteins those are responsible for the synthesis of proteins in the cell (30, 31), were downregulated in IL-8 treated samples, including *RPL39*, *RPS21*, *RPS28*, *RPS29* (Supplemental Figure 6, A-C). While some components that involve in mitochondrial respiratory chain, including *MT-ND1*, *MT-ND2*, *MT-ND4*, *MT-CO3*, *MT-CYB*, *MT-ATP8* (32–34) were upregulated (Supplemental Figure 5A). They comprise essential bioenergetic pathway powering fibroblasts proliferation and migration, to expedite healing (Supplemental Figure 6, B and C). It has been reported that *ZNF580* mediates H_2_O_2_-induced leukocyte chemotaxis by promoting the release of IL-8 (35). Implying a positive feedback loop might augmenting the effect. We also noted S100 calcium-binding protein A6 (*S100A6*) which has been associated with proliferation, invasion, migration and angiogenesis of cancer cells (36), was also downregulated, implying a setting dependent manifestation between tumor and wound healing events (Supplemental Figure 6A).

Given the prominence of transforming growth factor β (TGF-β) in fibroblast activation, we finally sought to determine whether IL-8-mediated effects is similar to those of TGF-β. To pinpoint this, we also performed the RNA-seq to probe the transcriptomes of fibroblasts in response to IL-8 or TGF-β. We found very distinct differences between them. Though IL-8 treated samples resulted in downregulated genes involved in fibrotic events, including extracellular matrix organizers (*COL4A1*, *COL10A1*) (37, 38), IL11 (39, 40). However, it can enhance canonical signaling drives fibrotic programs, as evidenced by upregulated *TGFβR3* and *SMAD3* that are essential in fibrosis (41, 42) (Supplemental Figure 6, D-F). Thus, our findings might implicate a profibrotic role for IL-8 upstream of TGF-β which need further exploration. Additionally, some biomarkers that have been implicated in fibrosis (43, 44), were also upregulated in IL-8 treated samples compared with TGF-β treated group when we check top candidates. For example, semaphorin 3A (*SEMA3A*) and phosphodiesterase-5A (*PDE5A*) (Supplemental Figure 6D). These results collectively underscore the profibrotic effects of IL-8 on cutaneous fibroblasts are differs from effects of TGF-β, whereas to some extent is parallel.

## DISCUSSION

In current study, we pay our attention to human unique skin fibrotic disease-keloids, though lacking relevant animal models, and therefore difficult to study in terms of etiology and pathogenic mechanisms. While some genetic and environmental factors associated with keloid formation are identified (5, 7–12), none of them can be considered as a main explanation. It is highly probable that the formation of keloids is indeed a resultant of multiple factors. We noticed that current researches on keloid pathology overlook the effects of skin microbiota, even though bacteria are recognized as critical components of skin biology. We asked if bacteria live in keloid scars and what happens if they do. Herein, we found that keloids indeed contain bacteria, which may accompany by the immune response. We identified IL-8 as the factor induced in the presence of bacteria, which support pathogenic programs in keloid scarring (Figure 6).

**Figure 6.**
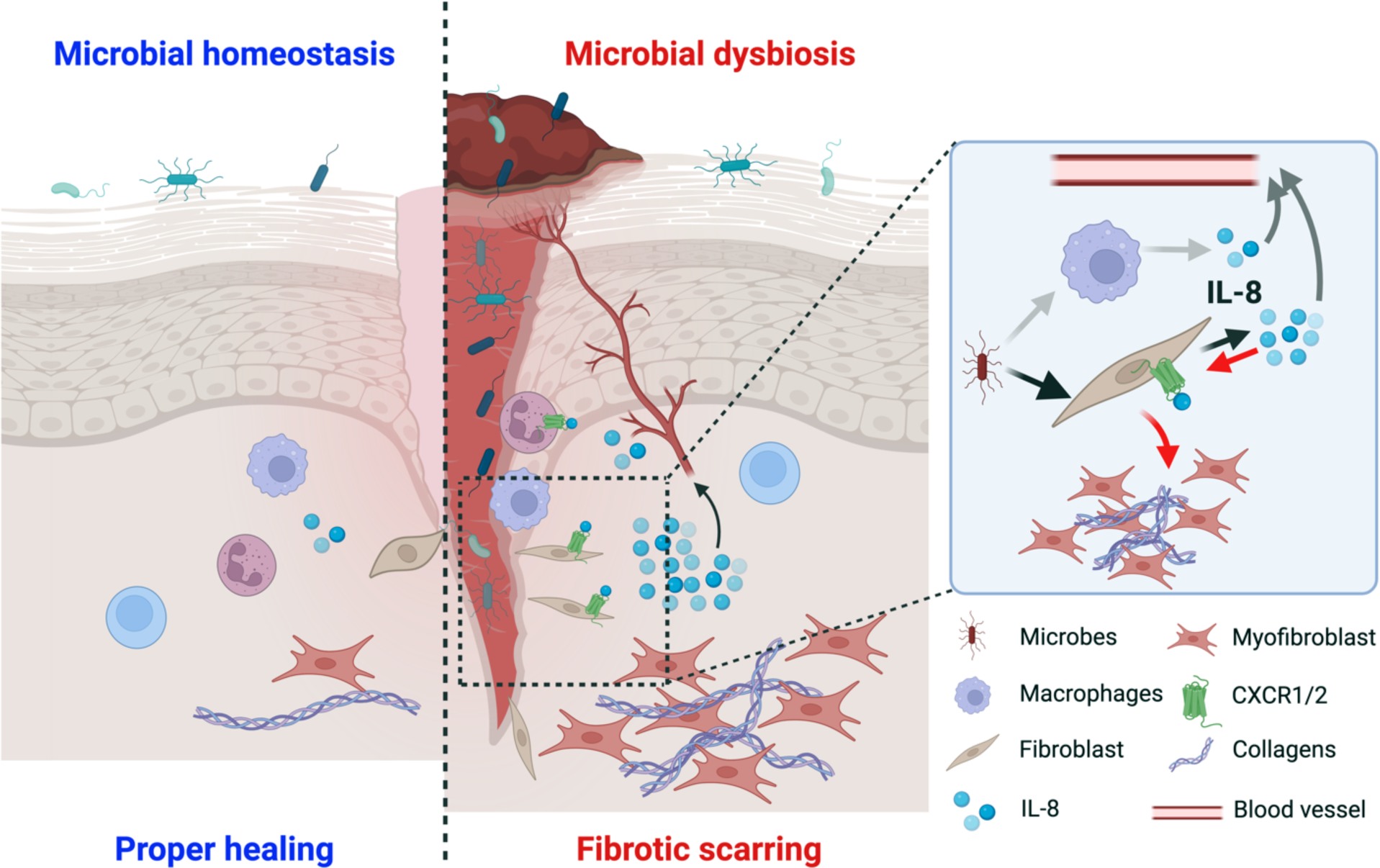
IL-8 and microbiota in keloids. Microbiota and IL-8 may play a role in development or sustaining of keloids. Bacteria present in the keloid scar induce production of IL-8, which in turn can bind to CXCR1/CXCR2 receptors on fibroblasts and stimulate collagen deposition. IL-8 is also released to circulation, but in this case its further role needs to be determined. The figure was created with BioRender.

Bacteria are traditionally seen as pathogens that need to be eliminated. Examples include typical immune-related mechanisms, like exfoliating the skin, producing anti-microbial peptides, and regulating the skin’s pH. Bacteria in wounds can be deadly, such as *Clostridium tetani* or *Clostridium perfringens*. However, microbiota can aid wound healing by affecting the skin immune system (45–47). From this point of view, bacteria can play a role of a double-edge sword, which can be both beneficial or detrimental for the outcome of wound healing. For example, *Staphylococcus epidermis* induces CD8^+^ T cells response and production of IFN-γ and IL-17, which protects against infection with other microorganisms (48) and support tissue repair (49). Staphylococcal lipoteichoic acids also activate TLR3 signaling, which reduces skin inflammation after wounding (50). However, skin injury is also a point of entry of microorganisms into deeper parts of the skin tissue (51). Finally, pathogenic bacteria play a critical role in pathological chronic wound healing, like diabetic wounds (52). Together, those studies show that the healing process depends on the bacteria in the wound.

Keloids may be considered as wounds that did not heal properly and the healing process is not yet finished. This agrees with the notion that keloid is a tumor-like disease, which shared likewise features with wound healing (53, 54). The healing process is usually divided into a few consecutive steps: inflammation, proliferation, and remodeling (53, 55). At the proliferation step, the many skin cell populations, including endothelial, epithelial and fibroblastic cells migrate into the wound and begin scar formation. Next, these processes are silenced and the tissue undergo remodeling to finish scar tissue maturation. In keloids, the inflammation and proliferation steps continue, and microbiota colonizing them may play a role. As we demonstrate with a tested panel of cytokines, keloid patients show increased concentration of proinflammatory cytokines in the serum, including IL-12 and IL-8. Interestingly, IL-8 is also a well-known pro-angiogenic factor (56). Presence of IL-8 in both the serum and tissue of keloid patients suggests ongoing proliferation and angiogenesis. Bacteria also induce IL-8 production by activation of TLR signaling in a variety of cells in the body (57–59). Here, we show that IL-8 is also produced after stimulation with heat-killed bacteria by skin fibroblasts. Therefore, skin cells’ secretion and cytokines like IL-8 may be regulated by microbiota in keloids, affecting keloid-associated fibroblasts.

This study has several restrictions. First, it was known that keloid patients often have a history of folliculitis or minor lesions, such as scratches on the skin surface. Bacteria can enter the skin through wounds and cause fibrosis by producing IL-8. The limited body volume of samples makes it challenging to determine if a particular bacterium is associated with keloids. This question requires further explorations with larger cohorts for a comprehensive understanding. Even more disturbing question is “hen and egg problem”. One symptom of keloid disease is itch, thus the bacteria present in the keloid tissue may be introduced after keloid formation by the patient. In such case, the bacteria within the tissue may be the effect, not the cause of the disease. Keloid tissue conditions can cause different bacteria to grow on the surface versus the inside. Also, our results did not show other microorganisms, like fungi, which has been showed that synergizing with bacteria in disease (60). This problem may be overcome by shotgun metagenomic analysis and ITS sequencing. Second, because of the human-specific character of keloids, we cannot use animal models and check the effect of bacteria in the in vivo setting. To this end, heat-killed bacteria is the way to stimulate in vitro, linking microbiota, fibroblasts and IL-8. Third, though our study elevated IL-8 as a critical mediator of bacteria herein, we cannot exclude other cytokines/inflammatory factors, perhaps even more important. Though we see the effects of IL-8 on fibroblasts, they are not as strong as expected. The experiments only focused on fibroblasts, without other vital cells like macrophages and neutrophils, which could partially explain the effect. Ideally, experiments using more sophisticated techniques (e.g., organoid etc.) which containing multicellular modules concurrently in the microenvironment would provide more comprehensive information, though it remains technically challenging to date. Therefore, we treat this study as a proof of the concept that microorganisms might be involved in the pathogenesis of keloids.

From a translational point of view, this study shows a new potential direction for clinical treatment of keloids. Currently, the most common therapeutic regimens are surgical excision, intradermal corticosteroid injection, laser or radiation therapy, pressure dressings, and combinations of them (1, 61, 62). For example, only approximately 50% of keloids respond to corticosteroid treatment, whereas nearly 100% regrow after surgery used as the sole therapy (62). Even with surgery and radiation therapy combined, 27% of our patients still had a relapse. None of these strategies targets microbiota, at least not directly. Antibiotics or phage treatments can improve outcomes for other therapies, like surgery, by reducing bacterial burden. One of the problems may be the high variability of bacteria identified in keloids. However, hi-throughput methods, like 16S rRNA sequencing, are now more affordable for diagnostic purposes. In such a case, keloids can be treated with personalized medicine using antibiotics that match the patient’s microbiome. Alternatively, identifying more molecular players between microbiota and keloids can lead to new target pathways. It is worthy to mention that growing evidence showed that IL-8 is such a molecule with prospective clinical value and deserves more attention (63–67).

In conclusion, our research shows that bacteria colonize keloids, and microbiota composition differs from normal skin. These microorganisms possibly induce the secretion of soluble factors, including IL-8, that further affect shared pathological events of fibrotic diseases evolution. Thus, our findings may stimulate further research on the bacteria and relevant pathways induced by their presence in keloids, which offer a promising avenue for translational practice.

## MATERIALS and METHODS

### Human subjects

Keloids were diagnosed on the basis of their history, anatomical location, clinical appearance, and pathology. Keloid (center) and margin tissues were harvested during plastic surgery from patients confirmed to have clinical evidence of keloid. Only mature keloids with no previous history of radiotherapy, chemotherapy, or intralesional steroids treatment prior to surgery were used in this research. Normal human upper eyelid skin samples were obtained from blepharoplasty of healthy donors and used for the experiments described in this study. The samples were acquired and processed by experienced pathologists.

### Cell lines and primary cultures

The adult human dermal fibroblast cell line (HDFα, FH1092, FuHeng Biology) was cultured in RPMI 1640 medium (11875093, Gibco) supplemented with 10% fetal bovine serum (FBS, 10099141C, Gibco) and 1% penicillin-streptamycin (10378016, Gibco). Cells were cultured at 37°C, 5% CO2 in a high-humidity atmosphere. The cells were authenticated by morphological examination and were routinely tested for the absence of mycoplasmic contamination. For isolation of primary fibroblasts, excised skin was immersed in physiological saline containing 2× antibiotics-antifungal agents (Anti-Anti, 15240062, Gibco), then immediately transferred to the laboratory. The skin tissue was washed twice in PBS containing 2× Anti-Anti. After removal of the adipose tissue under the reticular dermis, samples were cut into ≤5mm diameter pieces and incubated with dispase II (4942078001, Roche) for 2 h at 37°C. The epidermis was peeled off and discarded and the dermis was minced into small pieces and digested at 37°C for 1 h following 4°C for overnight using collagenase IV (17104019, Gibco). The resulting cell suspension was filtered through a 70 μm cell strainer (BD Falcon), and centrifuged at 400×g for 10 min. The supernatant was removed and the pellet was washed twice with PBS containing 2×Anti-Anti at 400×g for 5 min. The cells were then resuspended in DMEM medium (10569010, Gibco) supplemented with 10% FBS and 1% Anti-Anti (Gibco) for expanding.

### Immunohistochemistry staining

The tissues were fixed in 4% paraformaldehyde and paraffin-embedded. Tissues were cut in 8μm for further processing. For tissue morphology we used hematoxylin and eosin staining (H&E, 60524, Yeasen). For immunohistochemical (IHC) staining, antibodies against core part of lypopolisaccharide (HM6011, clone WN1 222-5, HycultBiotech, 1:500) and lipoteichoic acid (HM2048, clone 55, HycultBiotech, 1:200) were used according to standard procedures. Slides were scanned using the Aperio CS2 (Leica) automated slide scanner. The images were scored by the same observer following the standard pathohistological scoring method (0-4 score), and equal number of randomly selected visual fields (200× magnification) of each group were analyzed.

### mIHC staining

FFPE sections were evaluated by multiplex immunohistochemical analysis following previously reported protocol using antibodies specific for anti-α-SMA (ab124964, Abcam, 1:500), anti-IL-8/IL-8 (MAB208, R&D Systems, 1:100), anti-CD45 (2403794, Invitrogen, 1:200) and anti-CD68 (76437, Cell Signaling Technology, 1:400), cell nuclei were stained with 4,6-diamidino-2-phenylindole (DAPI, 62248, Thermo Scientific) (68). Slides were scanned by digital slide scanner and fluorescent images were obtained and analyzed with the confocal microscope and software (Nikon), and ImageJ software (National Institutes of Health, USA). For quantification of multiplex staining, the stained slides were scanned to obtain multispectral images using the Mantra System (PerkinElmer), which captures the fluorescent spectra at 20 nm wavelength intervals from 420 to 720 nm with identical exposure time. Images were deconvoluted and restitched. Reconstituted images underwent multiplex quantitative analysis using HALO image analysis software (Indica Labs). Density and percentages of signal or cell types were quantified.

### Immunofluorescence staining

Immunofluorescence staining was performed according to standard staining methods with deparaffinization and rehydration step, endogenous peroxidase quenching step, acidic antigen retrieval step, and blocking step prior antibodies’ hybridization. Primary antibodies (anti-FAP, 66562, Cell Signaling Technology, 1:100; anti-COLIA1 BA0325, BOSTER, 1:200 and anti-α-SMA, BM0002, BOSTER, 1:200, anti-Phalloidin C2203, Beyotime Technology, 1:100) were applied on slides at 4°C overnight and secondary antibodies (donkey anti-mouse IgG1 Antibody, Alexa Fluor 647, 34113, Yeasen, 1:200, Donkey anti-rabbit IgG Antibody, Alexa Fluor 488 34206, Yeasen, 1:200) were added for 1h at room temperature in the darkroom. Slides were mounted with DAPI Fluoromount-G™ anti-fluorescence quenching mountant (Yeasen, 36308). Slides were scanned using the automated slide scanner.

### 16s rRNA fluorescent in situ hybridization staining

FFPE tissues were used for 16S rRNA FISH staining following previously reported protocol (69). In brief, slides were deparaffinized, rehydrated and incubated in 70% ethanol at 4°C for 3 h. Slides were then washed in 2×saline sodium citrate buffer (SSC, AM9770, Invitrogen) and incubated with proteinase K (10 μg/mL in 2×SSC, AM2546, Invitrogen) for 15 min. After two washes with 2×SSC followed by two washes in 2×SSC with 15% formamide, slides were hybridized overnight at 30°C with Cy5 labelled probes (specific probe: GCTGCCTCCCGTAGGAGT [PMID: 2200342], non-specific control probe: CGACGGAGGGCATCCTCA, both at 1680 nM) in 2×SSC containing 10% dextran (60316, Yeasen), 1mg/mL *E. coli* tRNA, 0.02% bovine serum albumin (BSA, 36106, Yeasen), 2 mM vanadyl-ribonucleoside (S1402S, New England Biolabs), and 15% formamide (60345, Yeasen). Sections were washed for 20 min at 30°C in 2×SSC with 15% formamide. Cell nuclei were stained with DAPI in 2×SSC, 15% formamide at 30°C. Then slides were washed and mounted for scanning with the automated slide scanner (3D HISTECH).

### Microbiota collection, sequencing, and analysis

Surface or intra-tissue microbiota samples were collected from the pathological location or the normal lateral location of patients using a swab or punch biopsy (Catch-all Sample Collection Swab, Epicenter) moistened in Yeast Cell Lysis Buffer (from MasterPure Yeast DNA Purification Kit; Epicenter). Samples were snap frozen on dry ice, and DNA was isolated from specimens using the PureLink Genomic DNA Mini Kit (Invitrogen) and amplification of the 16S-V3+V4 region according to the manufacturer’s specifications. Sequencing of 16S rRNA amplicons was performed at the Penn Next Generation Sequencing Core using the Illumina Novaseq platform with 150 bp paired-end V3+V4 chemistry for the human samples.

Bioinformatic analysis was performed in R (70) v. 4.1.0. After initial trimming and filtering the raw data reads based on quality control, DADA2 algorithm (71) was used to obtain amplicon sequence variants (ASVs). After denoising and removal of chimeric sequences, the taxonomy was assigned based on SILVA v138 16S rRNA database (72) using DECIPHER package (73). Phylogenetic tree was constructed with phangorn (74). Sample data, taxonomy table, and phylogenetic tree were merge into phyloseq object and further analysed using tools from phyloseq package (75). We filtered out the phyla that were present in less than 3 samplesm agglomerated taxa at the genus level, and calculate relative abundances. Based on this, we calculated alpha and beta diversity. The analysis of minimum-spanning tree with Jaccard dissimilarity was performed with phyloseqGraphTest package (76). Krona charts were prepared using KronaTools (77). Functional pathways associated with microbiota were analysed using tax4Fun from themetagenomics package (78).

### Non-target metabolomic analysis

Non-target metabolomic analysis were performed using an Agilent Gas Chromatography Mass Spectrometer (7890A/5975C GC-MS System, Agilent, CA, USA) and conducted by Apexbio Co.

### Three-dimensional collagen gels

Primary human dermal fibroblasts suspended in serum-free medium were mixed with 4 mg/mL neutralized rat tail type I collagen at a ratio of 3:1. The mixture was seeded in 12-well plates at a cell density at 1×10^5^/mL. Gels were allowed to coagulation for 1 h at 37°C. The edge of the gel was detached from the well wall to allow contraction, and 1 mL culture medium (with/without 100 ng/mL rhIL-8) was added. Forty-eight hours later, the pictures were taken and the gel area was measured using ImageJ.

### Flow cytometry analysis

Adherent fibroblasts isolated from human that had labeled with 2 μM carboxyfluorescein succinimidyl ester (CFSE, C1031, Beyotime Biotechnology) were harvested and suspended in 500 μL FACS buffer. After maintaining in the dark and washed twice with FACS buffer, the cells were aquired by flow cytometry (BD FACSVerse). The data were analyzed using FlowJo software.

### Human dermal fibroblasts scratch assays

Cultured HDFα cells were allowed to reach confluency. Cells were pretreated with vehicle or 50 ng/mL recombinant human IL-8 (rhIL-8, R&D Systems) 12 h before scratch assay. Scratches were created by manual scraping of the cell monolayer with a 200-μL pipette tip. Cells were constantly treated with rhIL-8 (50 ng/mL). The scratch area was imaged using Nikon Ti2-U microscope and quantified using ImageJ software (n = 6).

### Trans-well assays

HDFα cells were evaluated for their ability to migrate a wound site through a trans-well assay. Cultured cells upon vehicle- or IL-8-treatment were seeded in the upper chamber of the trans-well plates and allowed migrating for 24h. Then those cells transferred into lower chamber were fixed in PFA for 30 minutes at room temperature (RT), stained with 0.4% trypan blue 30 min, washed 3×10 minutes with PBS, then dried in RT, photographed under 10×microscope (Ti2-U, Nikon) and cell numbers in the field were calculated manually by at least 2 colleagues (n = 6).

### RNA isolation and sequencing

RNA was extracted using RNeasy Mini Kit (Qiagen). RNA purity and concentration measurement, preparation of RNA library and transcriptome sequencing was conducted by TIANGEN Co. Genes with adjusted *p* value <0.05 and log2 (FoldChange) >0 were considered as differentially expressed. Heatmap was generated using R software.

### Immunoblotting

Proteins were extracted from cultured cells with RIPA lysis buffer (P0013C, Beyotime Biotechnology). Protein concentrations were measured using a coomassie brilliant blue G-250 bradford protein assay kit (P00006C, Beyotime Biotechnology). Protein extracts were separated by sodium dodecyl sulfate-polyacrylamide gel electrophoresis (SDS-PAGE) and electrophoretically transferred to nitrocellulose membranes (Millipore), and blocked in 5% nonfat dry milk. Blocking buffer was then removed, and the membranes were incubated with primary antibodies of α-SMA (19245, Cell Signaling Technologies, 1:1000), Col1A1 (BA0325, BOSTER, 1:2000) or GAPDH (30201, Yeasen, 1:2000) in Tris Buffered Saline Tween (TBST, 10mMTris-HCl, pH 7.5, 150 mM NaCl, 0.05% Tween 20) overnight at 4 °C, then washed 3×10 min with TBST, followed with horseradish peroxidaseconjugated anti-mouse or rabbit secondary antibody (Epizyme Technologies) for 1h at RT and washed with TBST 3×10 min. The membranes were incubated with ECL regents (Tanon) for the detection of the immunoreactive bands. Images were visualized using a Bio-Rad ChemiDoc Imaging System (Bio-Rad).

### Pathogens

Heat-killed *Salmonella typhimurium* (HKSA, tlrl-hkst2), and heat-killed *Staphylococcus aureus* (HKSA, tlrl-hksa) were purchased from InvivoGen. And the usage of them were following the manufacturer’s instruction.

### Clinical test of cytokines in plasma

Ethylenediaminetetraacetic acid (EDTA) anticoagulated blood from patients were collected at room temperature. The cytokine level from the plasma was quantitated using cytometric bead array kit (P110100403, JiangXi Cellgene) on Flow Cytometry (FASCantoII, BD). All measurements were handled according to the manufacturer instruction. The minimum and maximum limits of detection for all cytokines were 2.5 pg/mL and 2500 pg/mL, respectively. Cytokine concentrations were analyzed by FCAP Array v3.0.1 software and BD FACSDiva software.

### Enzyme-linked immunosorbent assay (ELISA)

Cultured HDFα cells were treated with diverse dosages of or without HKST or HKSA for 24h. Secreted IL-8 was measured using anti-human IL-8 ELISA kit (abs510004, Absin) according to the manufacturer’s instructions

### Statistical Analysis

Results represent means ± SEM. Data were analyzed using Prism 9.0 (GraphPad Software, San Diego, CA). Statistical significance was determined by an unpaired two-tailed Student’s t test when comparing two groups or one-way ANOVA when comparing more than two groups. Statistically significant differences were defined as ^ns^*p*>0.05, \**p*<0.05, \*\**p*< 0.01, \*\*\**p*<0.001 and \*\*\*\**p*<0.0001 and they were indicated in the figures and figure legends, where the number of independent biological replicates per group, and the number of independent experimental replicates were denoted when needed.

### Study approval

All samples were collected under ethical approval of the Medical and Ethics Committees of Renji Hospital, Shanghai Jiao Tong University, and each patient signed an informed consent before enrolling in this study.

### Data availability

All data needed to evaluate the conclusions in the paper are present in the paper or the Supplementary Materials. The underlying data from 16S rDNA sequencing and RNA-Seq are available from the corresponding authors TM or RC upon rational request.

### Author contributions

RC, TM and WZ contributed to the experimental design, data analysis, and writing of the manuscript. RC, QP, ZZ and ZF prepare and provide clinical resources. WZ, QP, XH, QH, BS, FL and NZ performed the investigation. WZ and TM performed the formal analysis, data curation and visualization. RC and TM supervised the project. WZ and TM wrote the original draft. RC, TM, WZ and QP reviewed and edited the manuscript, all other authors read the manuscript and give useful suggestions.

## Supporting information

Including in the file

## ACKNOWLEDGMENTS

We thank all members in the team for their kind help and useful discussion. This work was supported by National Sciences Foundation of China (NSFC) grant, 32270957, 8201101169 to Tomasz Maj, 31700762 to Rui Chen, 82001722 to Wenyu Zhang; China Postdoctoral Science Foundation grant 2020M671147 to Wenyu Zhang; Renji Hospital start-up funding to Tomasz Maj. Address correspondence to: Tomasz Maj, Institute of Molecular Medicine, Renji Hospital, School of Medicine, Shanghai Jiao Tong University, Pujian Road 160, Shanghai 200240, China. Phone: 86.021.68382291. Or to: Rui Chen, Department of Surgery, Division of Plastic and Reconstructive Surgery, Renji Hospital, School of Medicine, Shanghai Jiao Tong University, Pujian Road 160, Shanghai 200240, China. Phone: 86.021.68385611.

## Conflict of interest

The authors have declared that no conflict of interest exists.

## Notes

### Competing Interest Statement

The authors have declared no competing interest.

