## Supplementary material for "Commensal microbiome dysbiosis elicits IL-8 signaling to drive fibrotic skin disease": Including in the file

•

Wenyu Zhang *et al*.

**This file includes:**

Figs. S1 to S6

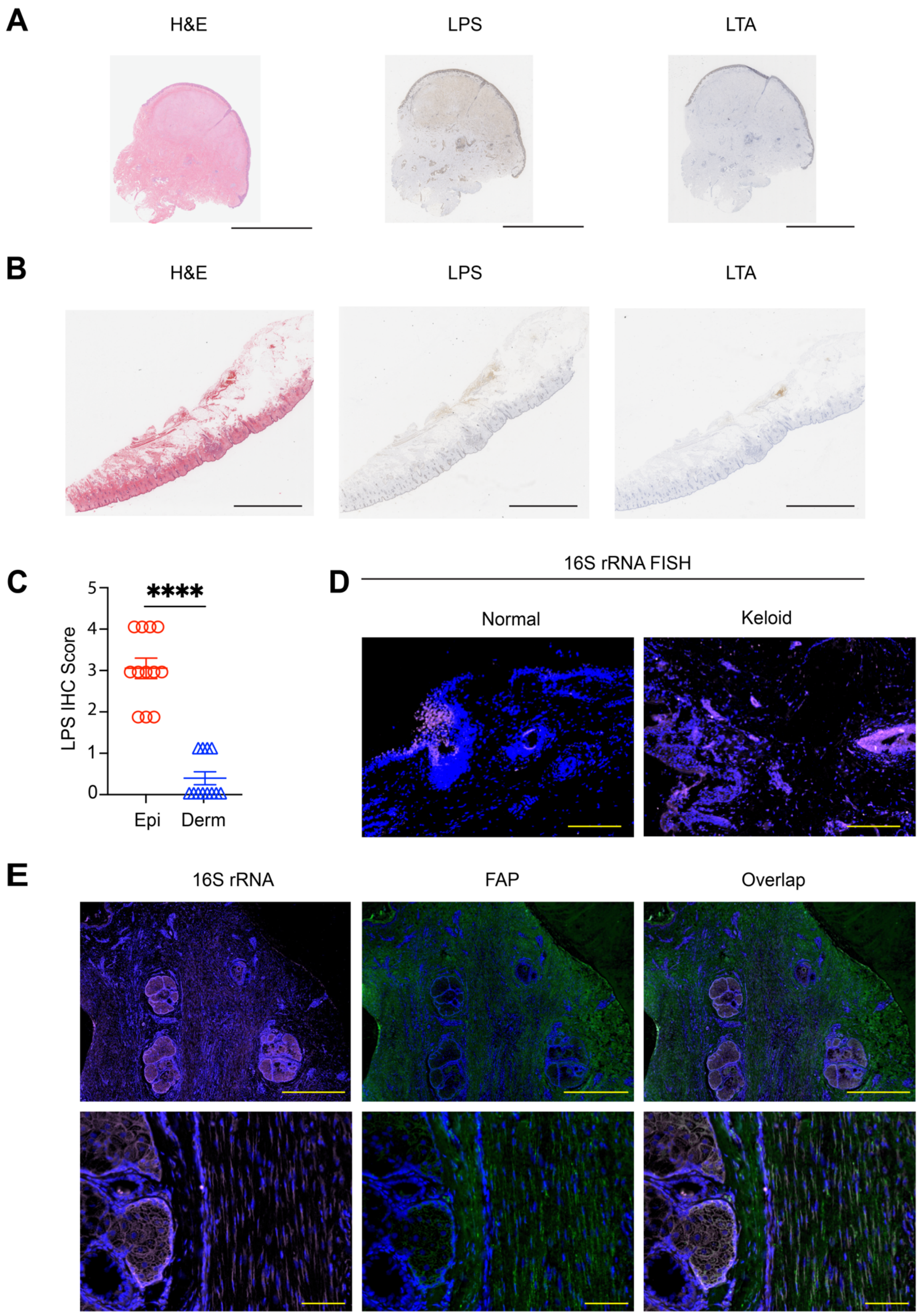

A
H&E
LPS
LTA
B
H&E
LPS
LTA
C
LPS IHC Score
5
4
3
2
1
0
****
Epi
Derm
D
16S rRNA FISH
Normal
Keloid
E
16S rRNA
FAP
Overlap

**Figure S1. Colonization of keloid by bacteria.**

(**A**) Global visualization of H&E and bacterial LPS, LTA in keloid tissues. n=12. Scale bars, 5000 μm. (**B**) Global visualization of H&E and bacterial LPS, LTA in normal tissues. n=2. Scale bars, 5000 μm. (**C**) Comparation of IHC scores of LPS and LTA staining in two layers of keloid skin (Epi and Derm, n=12). Mean ± SEM. ****$p < 0.0001$. Normality and Lognormality Tests followed by two-tailed paired Student's t test. (**D**) Normal and keloid tissues were subjected to 16S rRNA FISH, with the pan-bacteria EUB338 probe. Scale bars, 200 μm. (**E**) 16S rRNA FISH and FAP-IF staining on consecutive slides for representative keloid tissues are presented. n=6. Scale bars represent 1000 μm for low magnification images (top row) and 100 μm for high magnification images (bottom row).

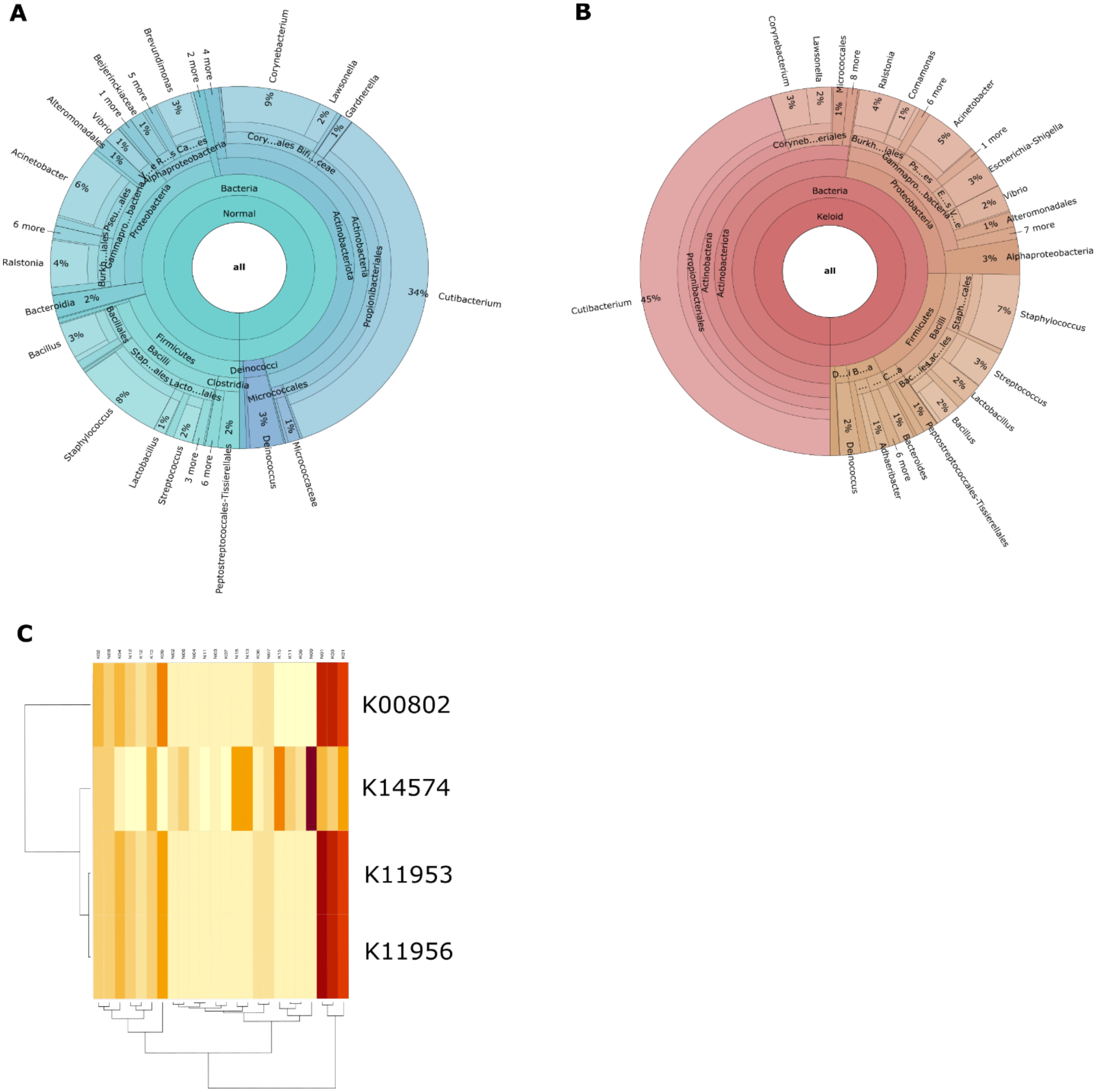


**Figure S2. Composition of bacteria in swab assay.**

(**A, B**) KRONA summary of bacterial communities analyzed by swab test on the surface of normal skin (**A**) and keloid scar (**B**). (**C**) Functional analysis of pathways distinctive for keloid and normal skin. The samples are shown in columns, where names beginning with N were aquired from normal skin, while samples with names beginning with K were collected from keloids.

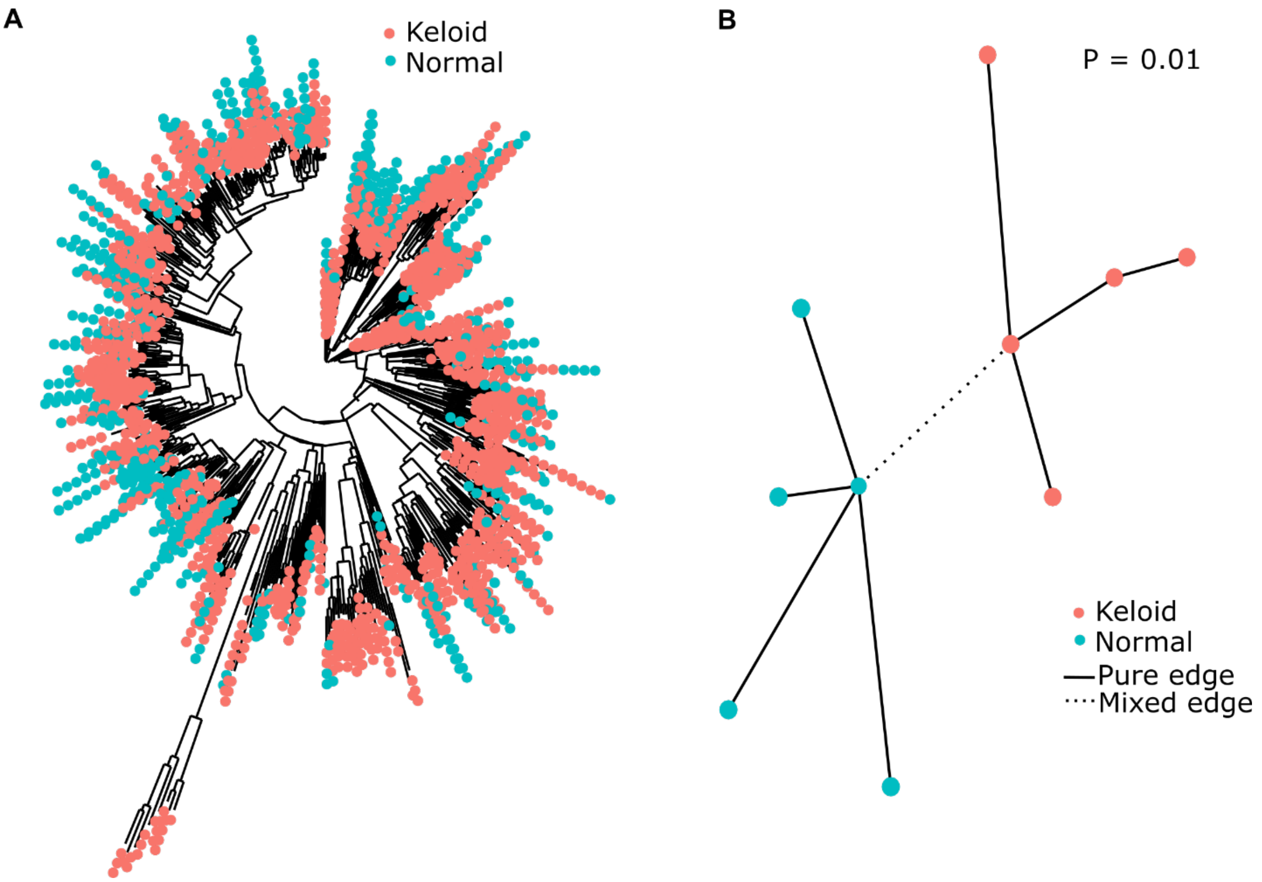


**Figure S3. Composition of bacteria inside normal skin and keloid tissue samples.**

(**A**) Phylogenetic tree of microbiota detected in the keloid and normal tissues. Each dot at the tip of tree means the presence of a given AVS in the collected material. E.g., three green dots and one red dot means that the AVS was present in three out of five normal skin samples, and two out of five keloid samples. (**B**) Permution test and tested network of bacterial communities in keloid and normal tissue samples.

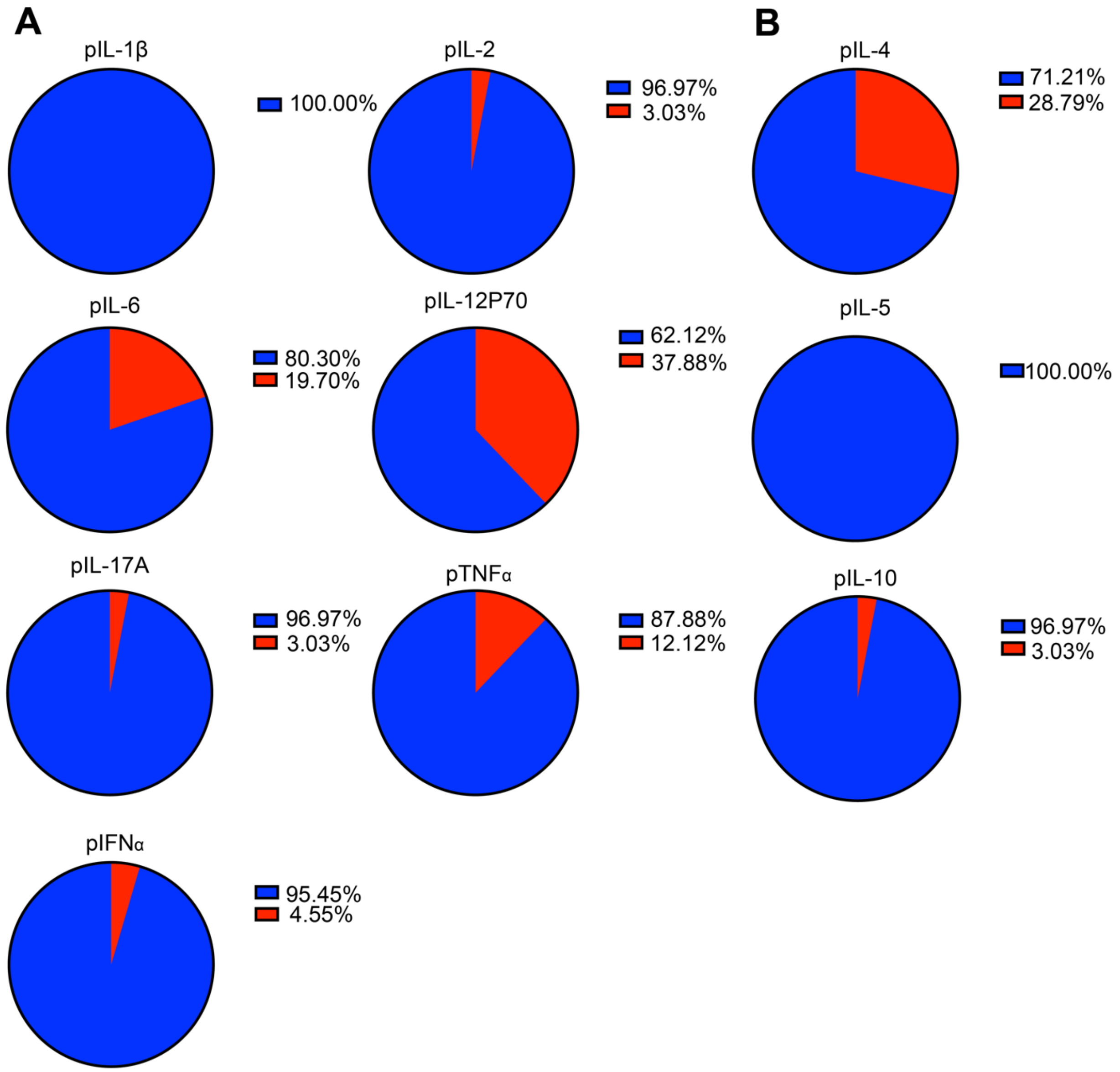
A
pIL-1β
100.00%
pIL-2
96.97%
3.03%
pIL-6
80.30%
19.70%
pIL-12P70
62.12%
37.88%
pIL-17A
96.97%
3.03%
pTNFα
87.88%
12.12%
pIFNα
95.45%
4.55%
B
pIL-4
71.21%
28.79%
pIL-5
100.00%
pIL-10
96.97%
3.03%

**Figure S4. Clinical tests for relevant cytokines from patients' blood samples.**

(**A**) Percentage of patients in tested inflammatory cytokine subgroup in circulation. Reference values（RVs） are as below: IL-1β=0~12.40 pg/ml, IL-2=0~5.71 pg/ml, IL-6=0~5.30 pg/ml, IL-12P70=0~3.40 pg/ml, IL-17A=0~20.60 pg/ml, TNFα=0~4.60 pg/ml, IFNα=0~7.42 pg/ml. (**B**) Percentage of patients in tested anti-inflammatory cytokine subgroup in circulation. RVs are as below: IL-4=0~3.00 pg/ml, IL-5=0~3.10 pg/ml, IL-10=0~4.91 pg/ml. Color in red indicates abnormal（above RVs）section. Color in blue indicates normal（within RVs）section. Total samples=66.

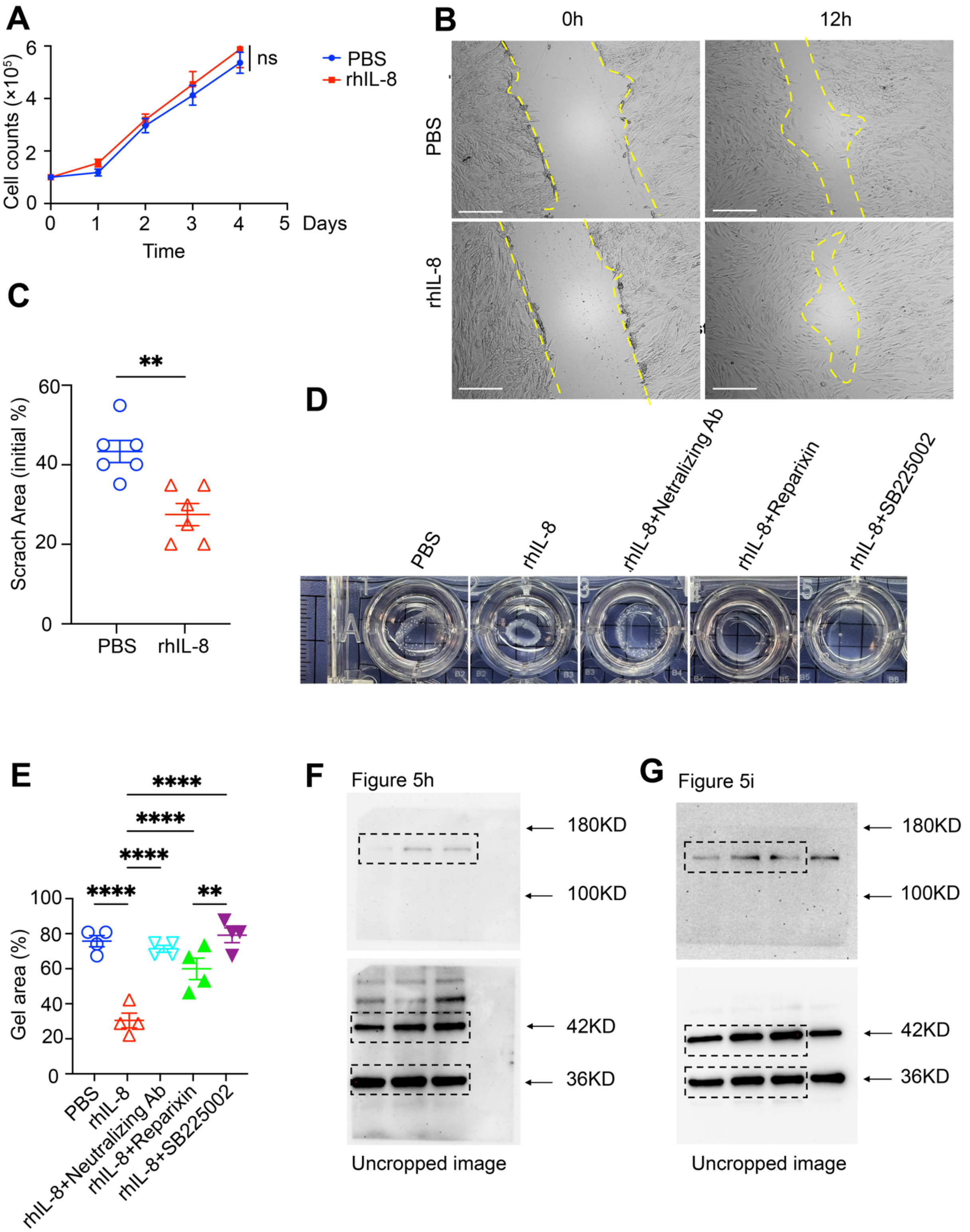
A
Cell counts (×10^5)
6
4
2
0
0
1
2
3
4
5
Days
Time
ns
PBS
rhIL-8
B
0h
12h
PBS
rhIL-8
C
**
60
40
20
0
Scrach Area (initial %)
PBS
rhIL-8
D
PBS
rhIL-8
rhIL-8+Netralizing Ab
rhIL-8+Reparixin
rhIL-8+SB225002
E
****
****
****
****
**
100
80
60
40
20
0
Gel area (%)
PBS
rhIL-8
rhIL-8+Neutralizing Ab
rhIL-8+Reparixin
rhIL-8+SB225002
F
Figure 5h
180KD
100KD
42KD
36KD
Uncropped image
G
Figure 5i
180KD
100KD
42KD
36KD
Uncropped image

**Figure S5. IL-8 mediates the changes in skin fibroblast behavior.**

(**A**) Cell Count Assays were executed by using automatic counter under indicated treatments: vehicle (PBS) and rhIL-8 (50 ng/ml). Data were collected at indicated time course and collectively showed as the growth curve (n=3). (**B**) Representative images showed rhIL-8 (50ng/ml) stimulation promotes the migration of primary human fibroblasts. Yellow dashed line marks scratch wound edges (n = 3 technical replicates, representative of two assays). (**C**) Scratched areas at 12h post wound are quantified as the percentage relative to the initial area at 0 hours. Scale bars, 100 mm. Mean ± SEM. **p < 0.01. Normality and Lognormality Tests followed by two-tailed unpaired Student's t test. (**D**) Gel contraction analysis of dermal fibroblasts with indicated treatment showed either neutralizing IL-8 (Anti-IL-8 antibody, 1μg/ml) or inhibition of rhIL-8 receptor (Reparixin or SB225002, 0.5μM for both) attenuate the promoted role of rhIL-8 (50ng/ml) in the activation of dermal fibroblasts. n = 4. (**E**) Gel area (%) were calculated and analyzed based on the data from (**D**). Mean ± SEM. **p < 0.01, ****p < 0.0001. Normality and Lognormality Tests followed by one-way ANOVA. (**F**) and (**G**) Uncropped blots related to Figure 5J and Figure 5K are shown, respectively.

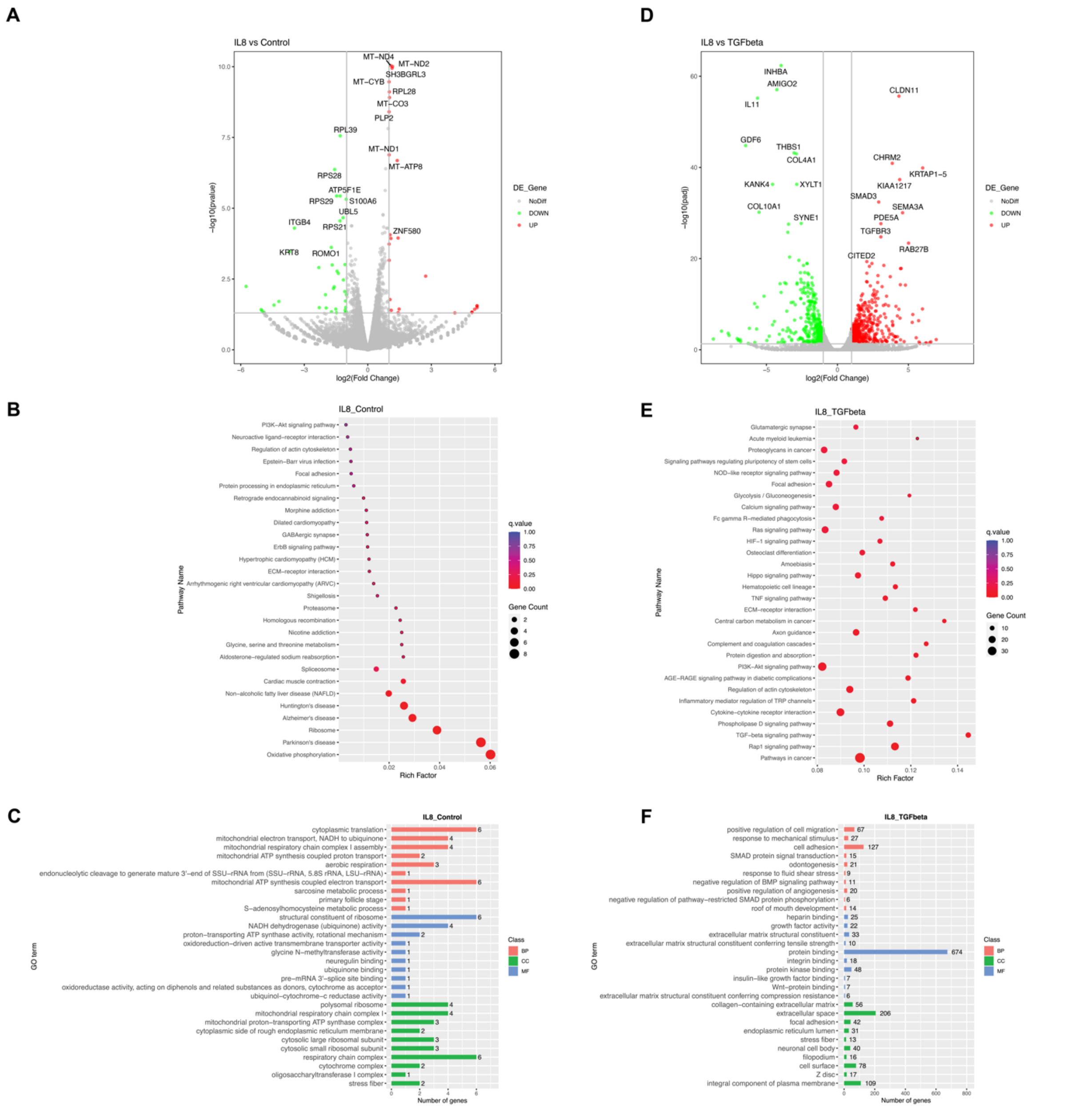


**Figure S6. Transcriptional shifts of the fibroblast upon IL-8 stimulation.**

(**A**) Differential expression analysis of response in freshly isolated dermal fibroblasts upon vehicle and rhIL-8 (50ng/ml). Significant upregulated genes (red) and downregulated genes (green) are shown, Top 10 genes are marked as indicated. (**B**) Gene Set Enrichment Analysis (GSEA) for genes from (**A**). (**C**) Gene Ontology（Go）analysis for genes from (**A**). (**D**) Differential expression analysis of response in freshly isolated dermal fibroblasts upon IL-8(50ng/ml) and TGFβ (10ng/ml). Significant upregulated genes (red) and downregulated genes (green) are shown, Top 10 genes are marked as indicated. (**E**) Gene Set Enrichment Analysis (GSEA) for genes from (**D**). (**F**) Gene Ontology（Go）analysis for genes from (**D**). Two independent experiments were executed on primary dermal fibroblast isolated from 2 patients.
